# A general method to fine-tune fluorophores for live-cell and *in vivo* imaging

**DOI:** 10.1101/127613

**Authors:** Jonathan B. Grimm, Anand K. Muthusamy, Yajie Liang, Timothy A. Brown, William C. Lemon, Ronak Patel, Rongwen Lu, John J. Macklin, Phillip J. Keller, Na Ji, Luke D. Lavis

## Abstract

Pushing the frontier of fluorescence microscopy requires the design of enhanced fluorophores with finely tuned properties. We recently discovered that incorporation of four-membered azetidine rings into classic fluorophore structures elicits substantial increases in brightness and photostability, resulting in the ‘Janelia Fluor’ (JF) series of dyes. Here, we refine and extend this strategy, showing that incorporation of 3-substituted azetidine groups allows rational tuning of the spectral and chemical properties with unprecedented precision. This strategy yields a palette of new fluorescent and fluorogenic labels with excitation ranging from blue to the far-red with utility in live cells, tissue, and animals.

## Introduction

Small molecule fluorophores are essential tools for biochemical and biological imaging^1,2^. The development of new labeling strategies^3^ and innovative microscopy techniques^4^ is driving the need for new fluorophores with specific properties. A particularly useful class of dyes is the rhodamines, first reported in 1887^5^, and now used extensively due to the superb brightness and excellent photostability of this fluorophore scaffold^1,2,6^. The photophysical and chemical properties of rhodamines can be modified through chemical substitution^6-13^, allowing the creation of fluorescent and fluorogenic labels, indicators, and stains in different colors^7,11-18^.

Despite a century of work on this dye class, the design and synthesis of new rhodamines remains severely limited by chemistry. The classic method of rhodamine synthesis—acid catalyzed condensation^2,5,6^—is incompatible with all but the simplest functional groups. To remedy this longstanding problem, our laboratory developed a strategy to synthesize rhodamine dyes using a Pd-catalyzed cross-coupling approach starting from simple fluorescein derivatives^19^. This approach facilitated the discovery of a novel class of dyes containing four-membered azetidine rings, which elicit substantial increases in the quantum yield relative to classic rhodamines containing *N*,*N*-dimethylamino groups^13^. Three first-generation azetidine dyes include the rhodamine-based ‘Janelia Fluor’ 549 (JF_549_, **1**, **Fig. 1a**), carborhodamine Janelia Fluor 608 (JF_608_, **2**), and Si-rhodamine Janelia Fluor 646 (JF_646_, **3**). The enhanced brightness and photostability of these dyes makes them exceptionally useful labels for single-molecule experiments *in vitro*^20^ and in living cells^13,21-24^.

**Figure 1.**
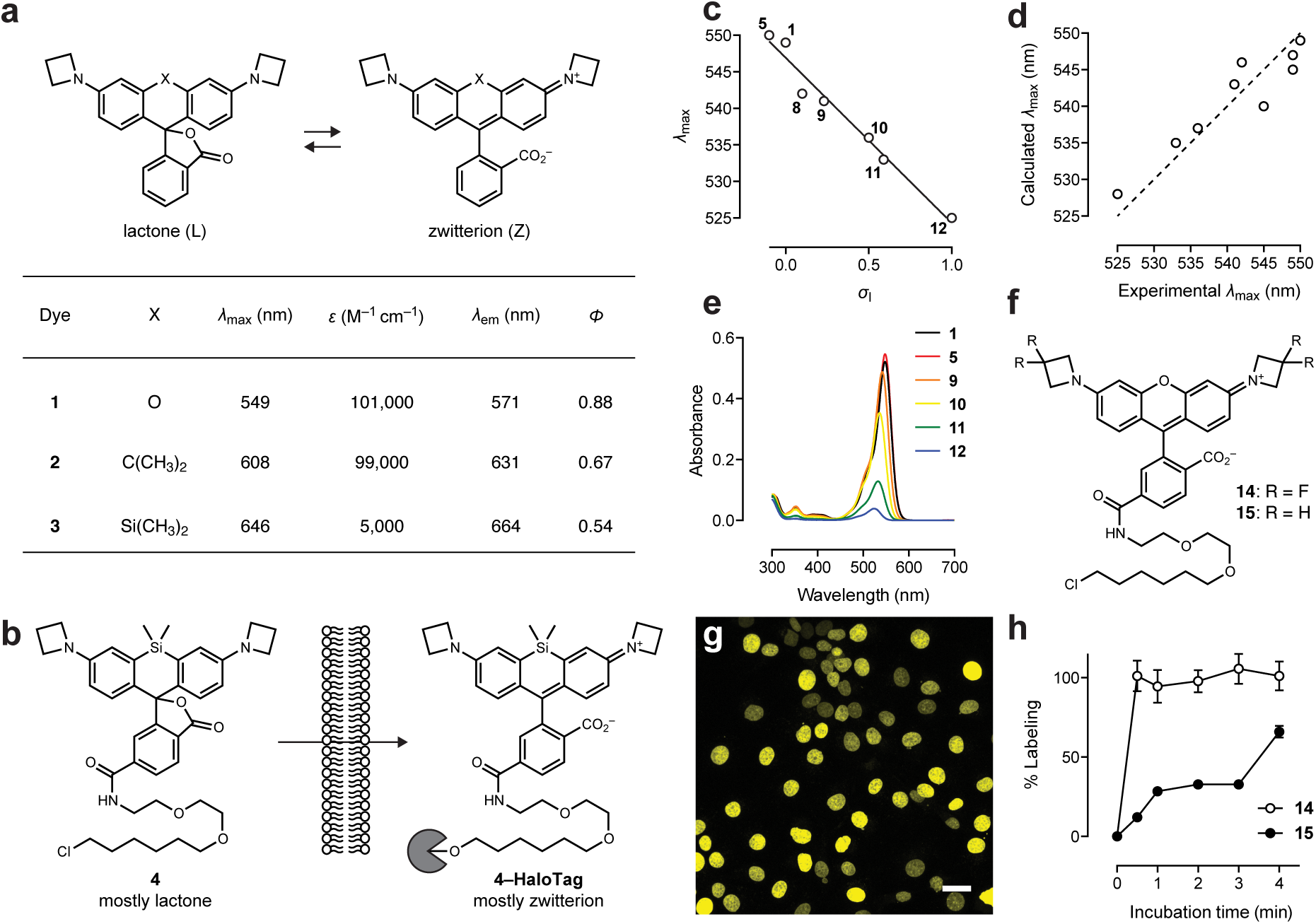
Fine-tuning rhodamine dyes using 3-substituted azetidines. (**a**) Properties of azetidinyl rhodamines **1–3**. All properties are taken in 10 mM HEPES, pH 7.3. (**b**) Structure of JF_646_–HaloTag ligand and schematic illustrating high cell permeability and fluorogenic labeling reaction. (**c**) Correlation of experimental λ_max_ *vs*. inductive Hammett constants (σ_I_ for dyes **1**, **5**, **8–12**. For the geminal disubstituted compounds **5** and **12** the σ_I_ of the substituent was doubled. Solid line shows linear regression (R^2^ = 0.97). (**d**) Correlation between calculated and experimental λ_max_ values for dyes **1**, **5–12**; dashed line shows ideal fit. (**e**) Absolute absorbance of **1**, **5**, **9–12** (5 μΜ) in 1:1 dioxane:H_2_O. (**f**) Chemical structure of JF_525_–HaloTag ligand **14** and JF_549_–HaloTag ligand **15**. (**g**) Image of live, washed COS7 cells expressing histone H2B–HaloTag fusions and labeled with ligand **14**. Scale bar: 30 μm. (**h**) Plot of percent labeling of histone H2B–HaloTag fusions in live cells vs. incubation time for ligands **14** and **15** determined by chasing with ligand **4**.

Although the Janelia Fluor dyes find broad use in cellular imaging^21-24^, it would be advantageous to develop methods to fine-tune the spectral and chemical properties of these dyes. The parent azetidinyl-rhodamine Janelia Fluor 549 (JF549, **1**; **Fig. 1a**) absorbs green light (λ_max_/λ_em_ = 549 nm/571 nm, *Φ* = 0.88) but is less suited for shorter excitation wavelengths. Replacing the xanthene oxygen in JF549 (**1**) with quaternary carbon yields JF_608_ (**2**) with a 59-nm shift in spectral properties (λ_max_/λ_em_ = 608 nm/631 nm, *Φ* = 0.67)—suboptimal for common orange excitation sources centered at 589 or 594 nm. Finally, inclusion of a silicon atom yields JF_646_ (**3**) with a 97-nm shift in spectra (λ_max_/λ_em_ = 646 nm/664 nm, *Φ* = 0.54). This dye is well matched for excitation with common far-red laser lines (*e.g*., 647 nm), although excitation with 633 nm light is less efficient. Overall, the ability to precisely adjust λ_max_ of rhodamine dyes could enable better matching of fluorophores with common excitation sources and filter sets.

In addition to λ_max_ and λ_em_, another critical property of rhodamine dyes is the equilibrium between the colorless, ‘closed’ lactone (L) form and the colored, fluorescent, ‘open’ zwitterionic (Z) form (**Fig.1a**). Both JF_549_ (**1**, *ε* = 1.01 × 10^5^ M^−1^cm^−1^) and JF_608_ (**2**, *ε* = 9.9 × 10^4^ M^−1^cm^−1^) primarily adopt the open form in water as evidenced by their high absorptivity. In contrast, JF_646_ (**3**) predominantly adopts the colorless lactone form in solution (*ε* = 5.0 × 10^3^ M^−1^cm^−1^). Likewise, the free JF_646_–HaloTag^25^ ligand (**4**) is primarily in the colorless, lipophilic lactone form in aqueous solution, which renders it highly cell permeable (**Fig. 1b**). The binding of HaloTag ligand **4** to its cognate protein shifts the L−Z equilibrium to the fluorescent form with a modest on:off absorbance ratio = 21 (**Fig.1b**)^13^. Thus, structural modifications that shift the L–Z equilibrium could be used to improve both the cell permeability and fluorogenicity of labeling with JF_646_ and other rhodamine fluorophores.

Here, we describe a general method to finely tune the spectral and chemical properties of rhodamine dyes with unprecedented precision. The use of 3-substituted azetidines on the Janelia Fluor scaffold allows modulation of λ_max_, λ_em_, and the L−Z equilibrium without affecting fluorescence quantum yield. The shifts in chemical and spectral properties can be explained using physical organic chemistry principles and quantum mechanical calculations. The structure–activity relationships are generalizable to rhodols, carborhodamines, and Si-rhodamine derivatives, allowing the rational design of improved fluorescent and fluorogenic labels across the visible spectrum.

## Results

### Synthesis of rhodamines with 3-substituted azetidine groups

We reasoned we could tune the physicochemical properties of JF_549_ (**1**) by exploring different substitution patterns on the azetidine ring. Indeed, the azetidinyl-rhodamine system provides an ideal test case for N-substituent effects due to the following: (1) the high-yielding Pd-catalyzed cross-coupling synthesis^19^, (2) the commercial availability of assorted 3-substituted azetidines, (3) the short, three-bond separation between the substituent and the rhodamine aniline nitrogen, and (4) the symmetry of the system. We hypothesized that electron withdrawing groups would decrease the λ_max_ of the fluorophore and also shift the L–Z equilibrium towards the closed, colorless form lactone based on reports of fluoroalkane-substituted rhodamine dyes^17,26^.

To test these predictions, we synthesized compounds **5**–**12** from fluorescein ditriflate (**13**) using Pd-catalyzed cross-coupling (**Table 1**). We then evaluated the photophysical properties of compounds **5**–**12** in aqueous solution, comparing them to JF_549_ (**1**; **Table 1**). All of the substituted azetidinyl dyes were synthesized in good yield and showed high **ε** values above 1 × 10^5^ M^−1^cm^−1^. The lone exception was the 3,3-difluoroazetidine compound **12**, which exhibited a slightly lower absorptivity (*ε* = 9.4 × 10^4^ M^−1^cm^−1^). Likewise, the quantum yield values of the azetidine dyes **5**–**12** were all >0.80 with the exception of the *N*,*N*-dimethyl-azetidin-3-amine compound **8**, which showed *Φ* = 0.57 at pH 7.4. The quantum yield value for **8** is rescued at pH 5.0 (*Φ* = 0.89; **Table 1**), suggesting photoinduced electron transfer (PeT) quenching by the unprotonated dimethylamino groups^27^.

**Table 1.** Synthesis and properties of azetidinyl rhodamines **1**, **5**–**12**.

| Dye | | Yield (%) | $\lambda_{\text{max}}$ (nm) | $\epsilon_{\text{H}_2\text{O}}$ ($\text{M}^{-1}\text{cm}^{-1}$) | $\epsilon_{\text{max}}$ ($\text{M}^{-1}\text{cm}^{-1}$) | $\lambda_{\text{em}}$ (nm) | $\Phi$ | $K_{\text{L-Z}}^c$ |
| --- | --- | --- | --- | --- | --- | --- | --- | --- |
| **1** |  | 95 | 549 | 101,000 | 134,000 | 571 | 0.88 | 3.5 |
| **5** |  | 86 | 550 | 110,000 | 143,000 | 572 | 0.83 | 3.2 |
| **6** |  | 54<sup>a</sup> | 549 | 111,000 | 138,000 | 572 | 0.87 | — |
| **7** |  | 47<sup>a</sup> | 545 | 108,000 | 130,000 | 568 | 0.87 | — |
| **8** |  | 80 | 542 | 111,000 | 127,000 | 565 | 0.57<sup>b</sup> | — |
| **9** |  | 83 | 541 | 109,000 | 137,000 | 564 | 0.88 | 2.5 |
| **10** |  | 89 | 536 | 113,000 | 141,000 | 560 | 0.87 | 1.0 |
| **11** |  | 85 | 533 | 108,000 | 133,000 | 557 | 0.89 | 0.24 |
| **12** |  | 91 | 525 | 94,000 | 122,000 | 549 | 0.91 | 0.068 |

<sup>a</sup>two-step yield after deprotection of the methyl ester intermediate
<sup>b</sup> $\Phi$ = 0.89 in pH 5.0 buffer
<sup>c</sup>equilibrium measured in 1:1 v/v dioxane:water

Although the *ε* and *Φ* of the different azetidinyl rhodamine dyes was largely immune to substitution at the 3-position, the λ_max_ and λ_em_ values were strongly affected by the nature of the substituent (**Table 1**). Groups with greater electron-withdrawing character elicited larger hypsochromic shifts in λ_max_. This effect was additive. For example, the 3-fluoroazetidinyl compound **10** showed a 13-nm blue shift (λ_max_ = 536 nm) relative to the parent dye **1** and the 3,3-difluoroazetidinyl-rhodamine (**12**) showed a further hypsochromic shift of 11 nm (λ_max_ = 525 nm). We plotted λ_max_ against the available Hammett inductive substituent constants (σ_I_)^28^ for the azetidine substituents in dyes **1**, **5**, and **8**–**12** and observed an excellent correlation (**Fig. 1c**), suggesting that the inductive effect of the substituents was primarily responsible for the decrease in absorption and emission maxima. The changes in λ_max_ were also observed in quantum mechanical calculations where the calculated λ_max_ values were within 5 nm of the experimental values (**Fig.1d**).

We then determined the *K*_L–Z_ for fluorophores **1**, **5**, and **9**–**12** to examine the effect of the azetidine substituents on the lactone–zwitterion equilibrium^29^; values for compounds **6**–**8** were not examined due to the ionizable substituents on the azetidine ring (**Table 1**). We determined these equilibrium values from the maximal extinction coefficients (ε_max_) measured in acidic media (Methods). JF_549_ (**1**) and the 3,3-dimethylazetidinyl-rhodamine 5 showed K_L–Z_ values >3, indicating that these dyes exist primarily in the open form. In contrast, the equilibrium of the 3,3-difluoroazetidinyl rhodamine (**12**) was substantially smaller (K_L–Z_ = 0.068), showing that the electron-withdrawing fluorine substituents can shift the equilibrium toward the closed lactone form. The remainder of the dyes exhibited K_L–Z_ values that were intermediate and trended with the electron-withdrawing character of the substituent (**Table 1**). Collectively, these results show that the λ_max_ and L–Z equilibrium can be rationally tuned using different 3-substituted azetidines without compromising fluorophore brightness.

### Janelia Fluor 525

Rhodamine **12** exhibits λ_max_ at 525 nm and a high quantum yield (*Φ* = 0.91), making it a useful label for imaging with blue-green excitation (514–532 nm). Based on the λ_max_, we named this fluorophore ‘Janelia Fluor 525’ (JF525) and prepared the JF525-HaloTag ligand (14, **Fig.1f**, **Supplementary Note**), which showed excellent labeling in live cells expressing HaloTag–histone H2B fusions (**Fig.1g**). We hypothesized that the JF_525_–HaloTag ligand (**14**) would show improved cell permeability relative to the analogous JF_549_–HaloTag ligand (**15**, **Fig.1f**) based on its higher propensity to adopt the lactone form (**Table 1**, **Fig.1e**, **Supplementary Fig.1a**). To test this hypothesis, we performed a pulse–chase experiment in live cells expressing HaloTag-histone H2B fusions pulsing with either JF_525_–HaloTag ligand 14 or JF_549_–HaloTag ligand **15** and chasing with the far-red JF_646_ ligand **4** (**Fig.1b**). Compound 14 labeled intracellular proteins faster than JF_549_ ligand **15 Fig. 1h**). These results support the hypothesis that shifting the L–Z equilibrium towards the lactone form can improve cell permeability and also validate JF_525_ as the first cell-permeable self-labeling tag ligand with an excitation maximum near 532 nm.

### Janelia Fluor 503

Since fluorine is the most electronegative atom, the difluoroazetidine-containing JF525 (**12**) represents the tuning limit of the azetidinyl-rhodamines towards the blue region of the spectrum. To access shorter wavelength dyes, we turned to the rhodol scaffold, which remains an intriguing but underexplored dye class^30^. In previous work^13^ we synthesized an azetindinyl-rhodol **16** (‘Janelia Fluor 519’), which exhibited λ_max_/λ_em_ = 519 nm/546 nm, *ε* = 5.9 × M^−1^cm^−1^, and *Φ* = 0.85 (**Fig.2a**). Based on the data with JF525 (**12**, **Table 1**) we surmised that replacement of the azetidine substituent with a 3,3-difluoroazetidine could elicit a desirable ∼15 nm blue-shift to yield a dye with maximal absorption closer to 488 nm.

**Figure 2.**
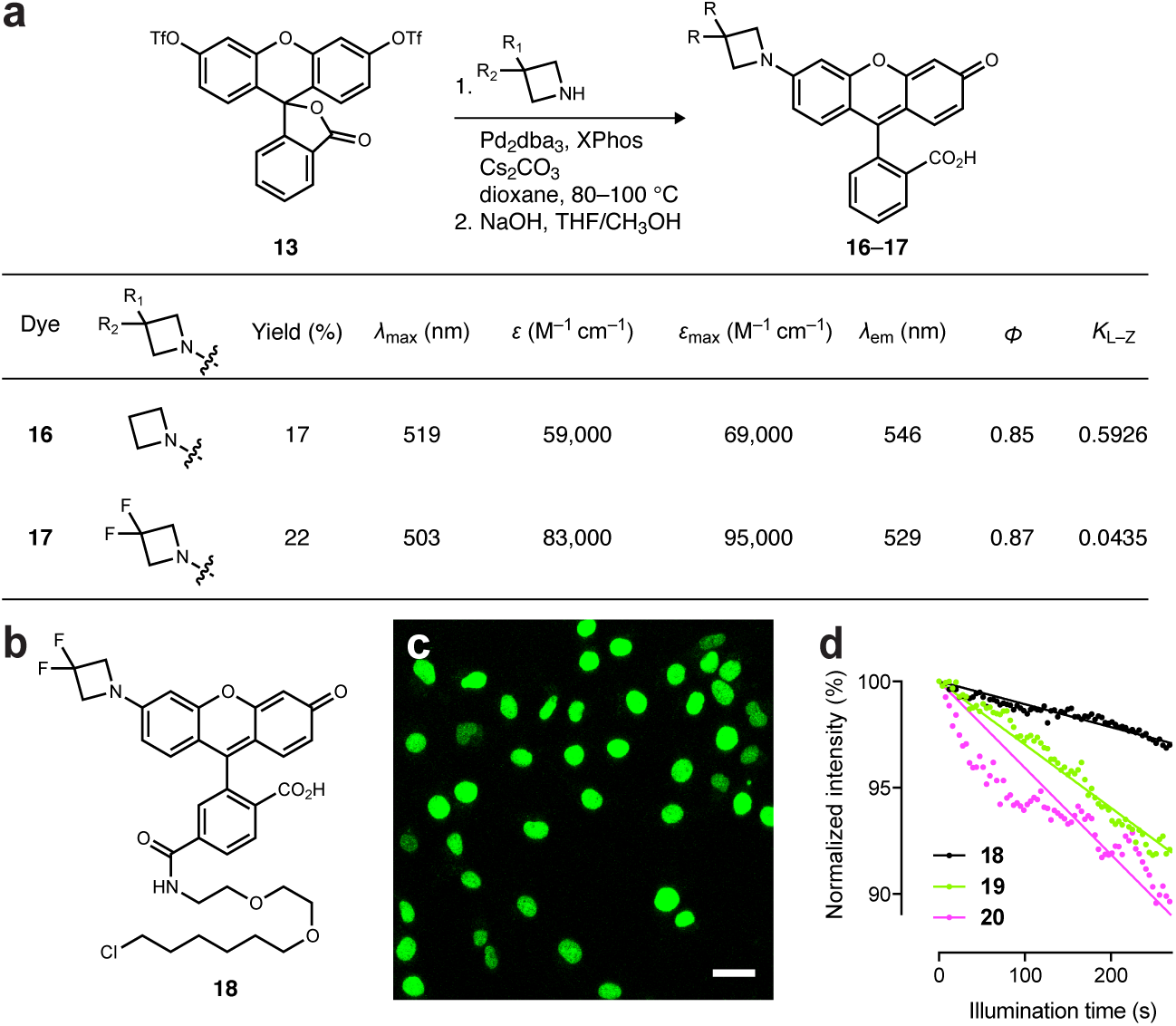
Design and synthesis of JF_503_. (**a**) Synthesis and properties of azetidinyl-rhodols **16−17**. Yields are for two steps; *K*_L–Z_ values were determined in 1:1 dioxane:H_2_O (**b**) Structure of JF_503_–HaloTag ligand **18**. (**c**) Image of live, washed COS7 cells expressing histone H2B–HaloTag fusions and labeled with ligand **18**. Scale bar: 35 μm. (**d**) Comparison of the photostability of cells labeled with 18 and cells labeled with 488 nm-excited dyes **19** and **20** (**Supplementary Fig.1b**); the initial photobleaching measurements are fitted to a linear regression.

To test this hypothesis, we synthesized the 3,3-difluoroazetidinyl-rhodol 25, which showed the expected blue-shifted spectra with λ_max_/λ_max_ = 503 nm/529 nm, *ε* = 8.3 × 10^4^ M^−1^cm^−1^, and *Φ* = 0.87 (**Fig. 2a**); we named this compound ‘Janelia Fluor 503’ (JF503). We then synthesized the JF503–HaloTag ligand (**18**, **Fig.2b**, Supplementary Note), which was an excellent label for histone H2B–HaloTag fusions live cells (Fig. 2c). We compared this novel label to two other 488-excitatable HaloTag ligands based on the classic rhodamine 110 (λ_max_/λ_em_ = 497 nm/520 nm; **19**, **Supplementary Fig.1b**) and the recently described *N*,*N*’-bis(2,2,2-trifluoroethyl)rhodamine (λ_max_/λ_em_ = 501 nm/525 nm; 20, **Supplementary Fig. 1b**)^17^. The cell-loading time course for all three dyes was similar (**Supplementary Fig.1c**) but the JF_503_ ligand showed higher photostability than the other two dyes in live cells (**Fig.2d**), consistent with previous reports comparing the photostability of rhodols to rhodamines^30^.

### Janelia Fluor 585

In previous work we synthesized several carborhodamine compounds from carbofluorescein ditriflate (**21**)^11^, including JF_608_ (**2**, **Fig.1a**, **Fig. 3a**)^13^. We also discovered that carborhodamines generally exhibit a higher propensity to adopt the closed lactone form, which was observed with JF_608_ (K_L–Z_ for **2** = 0.091). However, as noted above, JF_608_ exhibits a suboptimal λ_max_ and is not fluorogenic when used as a HaloTag label. We hypothesized that incorporation of 3,3-difluoroazetidine substituents into a carborhodamine structure could solve both problems. Based on the 3,3-difluoroazetidine compound **12** in the rhodamine series (**Table 1**), this modification was expected to elicit a blue-shift of approximately 24 nm, bringing the λ_max_ closer to the desired excitation wavelengths (e.g., 589 nm). We also predicted this modification would create a fluorogenic label, based on the higher propensity of carborhodamines to adopt the closed lactone form.^11^ We note previous efforts to shift the L–Z equilibrium of carborhodamines using direct halogenation on the xanthene ring system did produce a HaloTag ligand with modest fluorogenicity (on:off contrast = 9) but also severely decreased quantum yield^17^. Based on the data from the rhodamine series (**Table 1**), we expected that substitution with fluorine atoms on the 3-position of azetidine would *increase* quantum yield.

**Figure 3.**
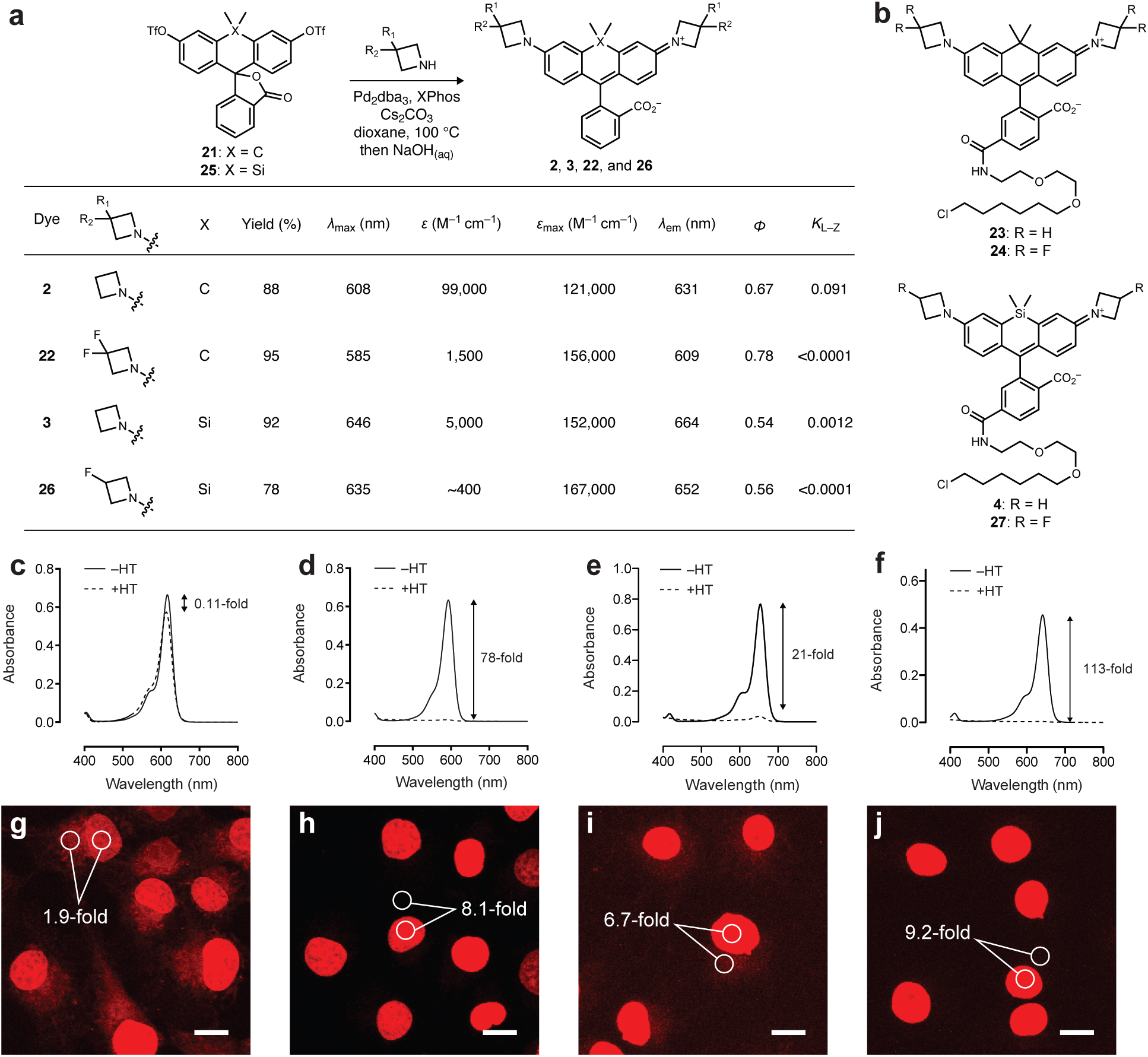
Design and synthesis of JF_585_ and JF_635_. (**a**) Synthesis and properties of azetidinyl-carborhodamines **2**, **22** and azetidinyl Si-rhodamines **3**, **26**. K_L–Z_ values were determined in 1:1 dioxane:H_2_O (**b**) Structure of HaloTag ligands derived from JF_608_ (**23**), JF_585_ (**24**), JF_64_**6** (**4**), and JF_635_ (**27**). (**c–f**) Absorbance of HaloTag ligands in the absence (−HT) or presence (+HT) of excess HaloTag protein. (**c**) JF_608_ ligand **23**. (**d**) JF_585_ ligand **24**. (**e**) JF_646_ ligand **4**. (**f**) JF_635_ ligand **24**. (**g–j**) Images of COS7 cells expressing HaloTag–histone H2B fusion and labeled with 250 nM of HaloTag ligands for **1** h and imaged directly without washing. The image for each dye pair was taken with identical microscope settings: λ_ex_ = 594 nm for g and h; λ_ex_ = 647 nm for **i** and **j**. Numbers indicate mean signal (**nuclear**) to background (cytosol; SNB) in three fields of view. (**g**) JF_608_ ligand **23** (SNB from n = 224 areas). (**h**) JF_585_ ligand **24** (SNB from n = 235 areas). (**i**) JF_646_ ligand **4** (SNB from n = 175 areas). (**j**) JF_635_ ligand **27** (SNB from n = 278 areas). Scale bars: 15 μm.

To test these predictions we synthesized the 3,3-difluoroazetidinyl carborhodamine (**22**) from 21 using Pd-catalyzed cross-coupling (**Fig.3a**). This dye showed the expected blue-shift in λ_max_ and λ_em_ showing 585 nm and 609 nm, respectively. The *Φ* was also modestly higher than **2** at 0.78, consistent with the results of the rhodamine series (**Table 1**). Based on these properties the dye was named ‘Janelia Fluor 585’ (JF_585_). However, unlike the 3,3-difluoroazetidine-substituted rhodamine analog **12** (JF_525_), which showed only a small decrease in *ε* relative to parent dye **1** (**Table 1**), carborhodamine **22** exhibited low visible absorption in water (*ε* = 1.5 × 10^3^ M^−1^cm^−1^) and a *K_L–Z_* near zero (Fig. 3a). These data show that inclusion of the electron-withdrawing fluorine substituents into this structure is sufficient to shift the equilibrium to the colorless lactone form. Measurement of the extinction coefficient in acidic trifluoroethanol, which stabilizes the open form of the dye, gave maximal extinction coefficient values ε_max_) of 1.21 × M^−1^cm^−1^ for fluorophore **2** and 1.56 × 10^5^ M^−1^cm^−1^ for fluorophore **22** (**Fig.3a**).

We prepared the JF_608_–HaloTag ligand (**23**) and JF585–HaloTag ligand (**24**, **Fig. 3b**) via Pd-catalyzed cross-coupling (**Supplementary Note**)^11,19^. To compare the utility and fluorogenicity of these new HaloTag ligands we first measured the absorbance of both **23** and **24** in the absence and presence of excess HaloTag protein. JF_608_–HaloTag ligand (**23**) showed only an 11% increase in absorption upon reaction with the HaloTag protein (**Fig.3c**). For JF585–HaloTag ligand **24**, however, the observed absorbance increase upon conjugation was substantially higher at 80-fold (Fig. 3d). This fluorogenic reaction enabled ‘no wash’ cellular imaging experiments: incubation of JF_608_–HaloTag ligand (**23**, 250 nM) with cells expressing HaloTag–H2B showed excellent nuclear labeling but high background due to the fluorescence of the free ligand staining internal membrane structures (Fig. 3g). In contrast, cells that were incubated with 250 nM of JF585–HaloTag ligand **24** and imaged directly showed bright nuclei with low fluorescence background (**Fig.3h**).

### Janelia Fluor 635

Having success with rational design of the carborhodamine system, we then turned to the Si-rhodamine JF_646_ (**3**, **Fig.3a**)^13^. Based on the rhodamine and carborhodamine data, we hypothesized that addition of a single fluorine atom on each azetidine ring would lower the absorptivity of the free dye, increase the chromogenicity of the labeling reaction, and cause a ∼13 nm hypochromic shift, thus attaining a λ_max_ near 633 nm. Accordingly, we converted Si-fluorescein ditriflate (**25**) to the fluorinated derivative **26** (**Fig.3a**), which showed λ_max_/λ_em_ = 635 nm/652 nm, a slightly higher *Φ* = 0.56 relative to JF_646_ (**3**), and an extremely low absorbance in water with an extinction coefficient value of approximately 400 M^−1^cm^−1^ (Fig. 3a). Based on these data we gave this dye the moniker ‘Janelia Fluor 635’ (JF_635_). Both JF_646_ (**3**) and JF_635_ (**26**) exhibit high extinction coefficient values in acidic media, with *ε*_max_ = 1.52 × 10^5^ M^−1^cm^−1^ and 1.67 × 10^5^ M^−1^cm^−1^, respectively (**Fig.3a**).

Analogous to the experiments with JF_608_ and JF_585_, we prepared the JF_635_–HaloTag ligand (**27**, **Supplementary Note**) and compared it to the JF_646_ ligand **4** (**Fig. 3b**). As mentioned previously, the JF_646_–HaloTag ligand (**4**) shows a 21-fold increase in absorbance upon binding to the HaloTag protein (**Fig 3e**). The shifted L–Z equilibrium observed for the free JF_635_ (**26**, **Fig.3a**) is also seen in the HaloTag ligand **27**, which shows exceptionally low background and a 113-fold increase in absorbance upon conjugation (**Fig.3f**). Both of these absorbance increases are much larger than the previously published SiTMR ligand **28**, which shows a 6.7-fold increase in absorption upon reaction with the HaloTag protein (**Supplementary Fig 1d–e**). These in vitro results were mirrored in no wash cellular imaging experiments, where we incubated 250 nM of either JF_646_ ligand 4 (**Fig.3i**), JF_635_ ligand 27 (**Fig.3j**), or SiTMR ligand 28 (**Supplementary Fig.1f**) with cells expressing histone H2B–HaloTag fusions. Cells labeled with either of the azetidinyl dyes (**4** or **27**) exhibited much lower nonspecific extranuclear fluorescence than the SiTMR compound **28**. However, as expected from the *in vitro* data (**Fig.3e–f**) the JF_635_ ligand **27** exhibited the highest contrast due to low fluorescence in the unbound state.

### Applications in tissue and *in vivo*

The HaloTag ligands of Janelia Fluor 585 (**24**) and Janelia Fluor 635 (**27**) are small, cell permeable, and exhibit high fluorogenicity upon labeling. We were curious if these properties would make them useful for labeling in more complex biological environments such as tissue or whole animals. We first attempted labeling in living brain tissue from *Drosophila* larvae using the JF_635_–HaloTag ligand (**27**) due to its far-red excitation (**Fig. 3a**) and high on:off ratio (**Fig.3f,j**). Nervous system explants from *Drosophila* third instar larvae expressing the HaloTag protein in ‘Basin’ neurons were dissected, incubated briefly with JF_635_–HaloTag ligand (**27**; 1 μM, 10 min), and imaged using the SiMView light-sheet microscope^31^. The JF_635_ label exhibited consistent labeling throughout the living tissue and low nonspecific background staining (**Fig.4a,b**), demonstrating its utility beyond simple cell culture.

**Figure 4.**
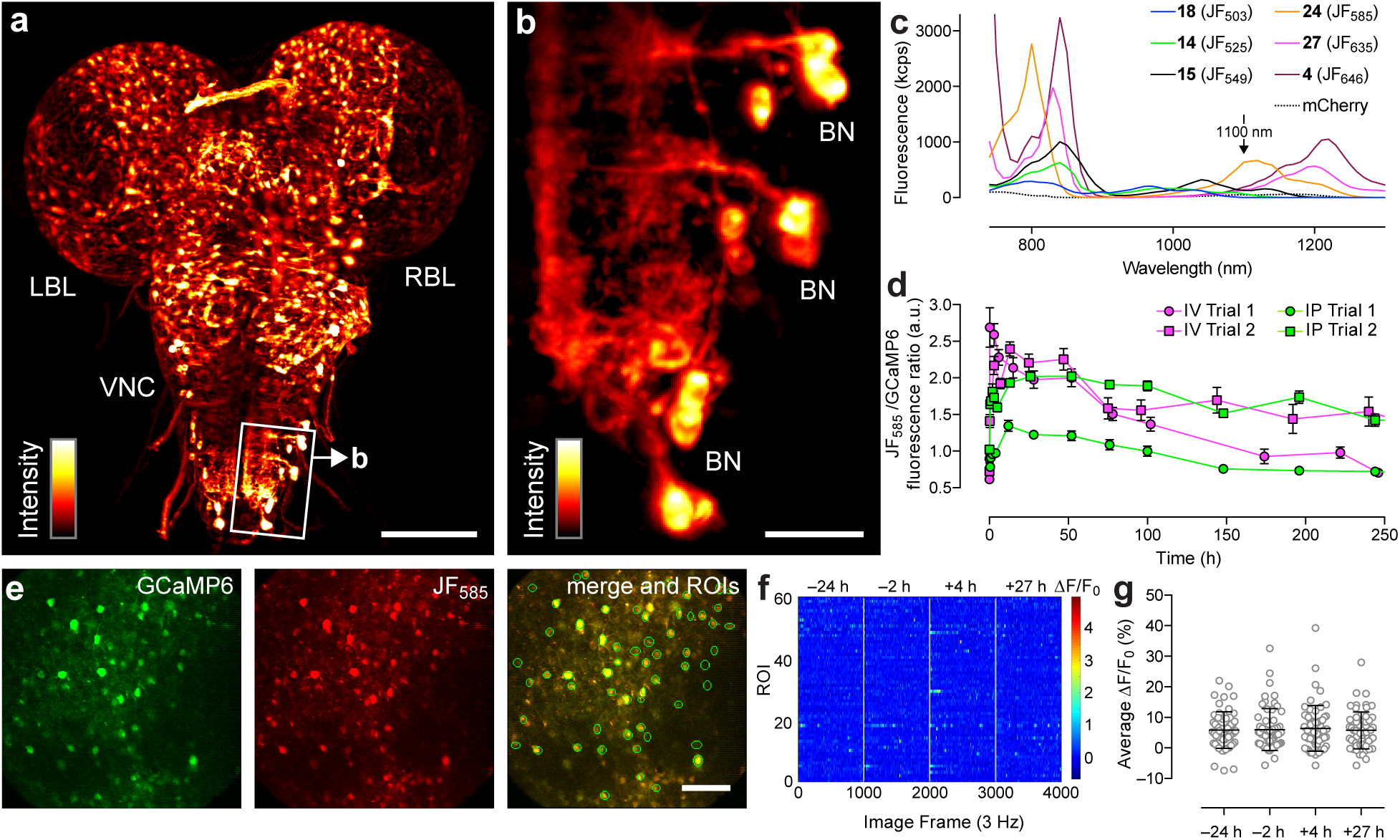
Labeling in tissue and *in vivo*. (**a**) SiMView light-sheet microscopy image of the central nervous system of a third instar *Drosophila* larva expressing myristoylated HaloTag protein in ‘Basin’ neurons (BNs) and stained with JF_635_-HaloTag ligand (**27**); LBL: left brain lobe; VNC: ventral nerve cord; RBL: right brain lobe. Scale bar: 100 μm. (**b**) Zoom in of boxed area in panel a showing individual BN cell bodies. Scale bar: 20 μm. (**c**) Two-photon fluorescence excitation spectra of HaloTag conjugates (**1** μM) from HaloTag ligands **4**, **14**, **15**, **18**, **24**, and **27** in 10 mM HEPES buffer (pH 7.3). The two-photon excitation spectra for mCherry is shown for reference. (**d**) Ratio of JF_585_ fluorescence to GCaMP6 epifluorescence at different time points after a single injection of JF_585_–HaloTag ligand (**24**, 100 nmol) either intravenous (IV) or intraperitoneal (IP); n = 2; error bars show ± s.e.m. (**e**) Two-photon microscopy images of neurons in layer 5 of the visual cortex coexpressing GCaMP6s (green) and JF_585_-labeled HaloTag (red) after IV injection of ligand **24** (*t* = 5 h). Scale bar: 100 μm. Green circles in the merged image indicate individual neurons as regions of interest (ROIs). (**f**) Raster plot of spontaneous neuronal activity in different ROIs (n = 61) before and after labeling with JF_585_–HaloTag ligand (**24**). (**g**) Plot of average spontaneous neural activity in each ROI before and after labeling with JF_585_–HaloTag ligand (**24**); error bars show ± s.d.; no significant difference is observed between time points (one-way ANOVA: p = 0.95).

We next evaluated the JF dyes in the brains of living mice. The JF_585_–HaloTag ligand **24** was chosen based on its high fluorogenicity (**Fig.3d,h**) and superior 2-photon fluorescence at 1100 nm excitation (**Fig.4c**), which is sufficiently separated from GFP-based indicators such as GCaMP6 (2-photon λ_max_ = 940 nm)^32^ to allow multicolor imaging. Cytosolic HaloTag protein was co-expressed in layer 4 or layer 5 visual cortex (V1) neurons with GCaMP6s via viral transduction and the mice were fitted with a chronic cranial window (**Methods**). Injection of 100 nmol of HaloTag ligand **24** into the tail vein (intravenous, IV) showed the JF_585_ ligand was blood-brain barrier-permeable and gave measureable labeling in the brain within 5 minutes, peaking around 6 hours and lasting for nearly two weeks as measured by epifluorescence (**Fig.4d**, **Supplementary Fig. 2a**). Intraperitoneal injection (IP) also showed effective delivery to the brain although the peak intensity was lower than IV administration (**Fig. 4d**). Under 2-photon imaging, we observed that the GCaMP6 and JF_585_ signals co-localized (**Fig.4e**, **Supplementary Video 1**) and the labeling showed no significant effect on neuronal activity (**Fig.4f,g**), establishing the utility of this fluorophore *in vivo*.

## Discussion

Replacement of *N*,*N*-dialkyl groups in rhodamine dyes with azetidines greatly improves the quantum efficiency without greatly affecting other spectral and chemical properties. Here, we show the azetidine strategy is remarkably flexible—substitutions on the azetidine ring do not compromise the high quantum yield values of the Janelia Fluor dyes and allow fine-tuning of λ_max_/λ_em_ and the L–Z equilibrium. This resulted in the development of JF_525_ (**12**) and its HaloTag ligand **14**—the first ligand for self-labeling tags with absorption maximum near 532 nm. The design principles we discovered are generalizable to analogous structures—rhodols, carborhodamines, and Si-rhodamines—allowing the rational design of finely-tuned fluorophores such as JF_503_ (**17**, **Fig. 2a**), JF_585_ (**22**), and JF_635_ (**26**, **Fig.3a**). Together with JF_549_ (**1**) and JF_646_ (**2**, **Fig.1**)^13^, we have now described six dyes that span the visible region of the spectrum and match common excitation wavelengths for fluorescence microscopy. In addition to the HaloTag ligands described above, we also prepared the corresponding SNAP-tag ligands **29**–**32**, which can label SNAP-tag fusion proteins inside live cells (**Supplementary Fig.2b–i**). Importantly, the new HaloTag ligands derived from JF_585_ and JF_635_ show a high degree of chromogenicity and fluorogenicity (**Fig.3c–j**), a critical parameter in advanced imaging experiments^33^. Of particular interest is the ability to deliver these two dyes to neural tissue in explants (**Fig. 4a,b**) or whole animals (**Fig.4d–g**), which could allow the imaging of deeper structures in the brain or the *in vivo* assembly of semisynthetic indicators for monitoring cellular activity^3^.

Although we focused on the fluorine-substituted azetidines in this paper, the other substitutions (**Table 1**) could be exploited to prepare fluorophores for specific applications. For example, the carboxy groups in compounds **5** and **6** could serve as attachment sites for a variety of chemical modifiers to improve solubility^7^, quench unwanted triplet states^34^, or allow the molecule to serve as a multivalent fluorescent cross-linker. The modest pH sensitivity and presence of the basic amine in compound **8** could allow it to function as a pH sensor or stain for lysosomes. The methoxy group on compound **9** could be elaborated to a polyethylene glycol (PEG) or other solubilizing group. Finally, the cyano group in compound **11** could be used in multimodal imaging regimes where both fluorescence and Raman^35^ modalities are used for imaging. In all, this general method to tune photophysical and chemical properties against a backdrop of high quantum yield will allow the precise design of many new fluorophores for specific, sophisticated biological imaging experiments in increasingly complex systems.

## Acknowledgements

We thank A. Berro and E. Schreiter (Janelia) for purified HaloTag protein, H. Choi (Janelia) for contributive discussions and a critical reading of the manuscript. This work was supported by the Howard Hughes Medical Institute.

## Author Contributions

L.D.L. and J.B.G. conceived the project. J.B.G. and A.K.M. performed organic synthesis. J.B.G. and L.D.L performed 1-photon spectroscopy measurements. Y.J., R.L. and N.J. designed, performed, and analyzed mouse imaging experiments. W.C.L. and P.J.K. designed, performed, and analyzed larval explant imaging experiments. R.P. and J.J.M performed 1-photon spectroscopy measurements. L.D.L. wrote the paper with input from the other authors.

## Online Methods

### Chemical synthesis

Methods for chemical synthesis and full characterization of all novel compounds can be found in the Supplementary Note.

### UV–vis and fluorescence spectroscopy

Fluorescent and fluorogenic molecules for spectroscopy were prepared as stock solutions in DMSO and diluted such that the DMSO concentration did not exceed 1% v/v. Spectroscopy was performed using 1-cm path length, 3.5-mL quartz cuvettes or 1-cm path length, 1.0-mL quartz microcuvettes from Starna Cells. All measurements were taken at ambient temperature (22 ± 2 °C). Absorption spectra were recorded on a Cary Model 100 spectrometer (Agilent). Fluorescence spectra were recorded on a Cary Eclipse fluorometer (Varian). Maximum absorption wavelength (λ_max_), extinction coefficient (*ε*), and maximum emission wavelength (λ_em_) were taken in 10 mM HEPES, pH 7.3 buffer unless otherwise noted; reported values for *ε* are averages (n = 3). Normalized spectra are shown for clarity.

### Determination *K*_L–Z_ and *ε*_max_

To determine *K*_L–Z_ we first performed dioxane–H_2_O titrations in spectral grade dioxane (Aldrich) and milliQ H_2_O (**Supplementary Fig. 1a**). The solvent mixtures contained 0.01% v/v triethylamine to ensure the rhodamine dyes were in the zwitterionic form. The absorbance values at λ_max_ were measured on 5 μM samples (n = 2) using a quartz 96-well microplate (Hellma) and a FlexStation3 microplate reader (Molecular Devices). Values of dielectric constant (*ε*_r_ were as previously reported^36^. We then calculated *K*_L−Z_ using the following equation^29^: *K*_L−Z_ = *ε*_dw_/*ε*_max_)/(1 − *ε*_dw_/*ε*_max_). *ε*_dw_ is the extinction coefficient of the dyes in a 1:1 v/v dioxane:water solvent mixture (**Fig.1e**); this dioxane–water mixture was chosen to give the maximum spread of *K*_L−Z_ values (**Supplementary Fig.1a**). *ε*_max_ is the maximal extinction coefficients measured in different solvent mixtures depending on dye type: 0.1% v/v trifluoroacetic acid (TFA) in 3,3,3-trifluoroethanol (TFE) for the rhodamines (**1**, **5**–**12**) and carborhodamines (**2**, **22**); 0.1% v/v TFA in ethanol for the Si-rhodamines (**3**, **26**); 0.01% v/v Et_3_N in TFE for the rhodols (**16**, **17**).

### Quantum yield determination

All reported absolute fluorescence quantum yield values (Φ) were measured in our laboratory under identical conditions using a Quantaurus-QY spectrometer (model C11374, Hamamatsu). This instrument uses an integrating sphere to determine photons absorbed and emitted by a sample. Measurements were carried out using dilute samples (*A* < 0.1) and self-absorption corrections^37^ were performed using the instrument software. Reported values are averages (n = 3). The quantum yield for compound **8** at pH 5.0 was taken in 10 mM sodium citrate buffer containing 150 mM NaCl.

### Molecular modeling

Computational experiments were performed using Gaussian 09.^38^ DFT and TD-DFT methods were used to calculate the spectral properties of the azetidinyl rhodamine compounds (**Table 1**). Calculations were performed at the B3LYP/6-31+G(d,p)/IEFPCM and TD-B3LYP/6-31+G(d,p)/IEFPCM theory levels for the ground states and excited states respectively. Frequency calculations confirmed that an energy minimum was found in geometry optimizations. Linear response solvation with the IEFPCM model was sufficient to study the excited state energies. Evaluations of TD-DFT theory have discussed the overestimation of excitation energies^39,40^, and previous studies of rhodamine excited states have reported using ∼0.4 eV correction to account for this overestimation^41,42^. We used a consistent –0.4 eV correction was applied to the calculated excited state energies, which gave good agreement with spectroscopy experiments (**Fig.1d**).

### Measurement of increase in absorbance of HaloTag ligands 4, 23, 24, 27, and 28 upon attachment with HaloTag protein

HaloTag protein used as a 100 μM solution in 75 mM NaCl, 50 mM TRIS·HCl, pH 7.4 with 50% v/v glycerol (TBS–glycerol). Absorbance measurements were performed in 1 mL quartz cuvettes. HaloTag ligands **4**, **23**, **24**, **27**, and **28** (5 μM) were dissolved in 10 mM HEPES, pH 7.3 containing 0.1 mgmL^−1^ CHAPS. An aliquot of HaloTag protein (1.5 equiv) or an equivalent volume of TBS–glycerol blank was added and the resulting mixture was incubated until consistent absorbance signal was observed (∼60 min). Absorbance scans are averages (n = 2).

### Multiphoton spectroscopy

HaloTag ligands **4**, **14**, **15**, **18**, **24**, and **27** (5 μM) were incubated with excess purified HaloTag protein (1.5 equiv) in 10 mM HEPES, pH 7.3 containing 0.1 mg·mL^−1^ CHAPS as above and incubated for 24 h at 4 °C. These solutions were then diluted to 1 μM in 10 mM HEPES buffer, pH 7.3 and the two-photon excitation spectra were measured as previously described^43,44^. Briefly, measurements were taken on an inverted microscope (IX81, Olympus) equipped with a 60×, 1.2NA water objective (Olympus). Dye–protein samples were excited with pulses from an 80 MHz Ti-Sapphire laser (Chameleon Ultra II, Coherent) for 710-1080 nm and with an OPO (Chameleon Compact OPO, Coherent) for 1000-1300 nm. Fluorescence collected by the objective was passed through a dichroic filter (675DCSXR, Omega) and a short pass filter (720SP, Semrock) and detected by a fiber-coupled Avalanche Photodiode (SPCM_AQRH-14, Perkin Elmer). For reference, a two-photon excitation spectrum was also obtained for the red fluorescent protein mCherry (1 μM), in the same HEPES buffer. All excitation spectra are corrected for the wavelength-dependent transmission of the dichroic and band-pass filters, and quantum efficiency of the detector.

### General cell culture and fluorescence microscopy

COS7 cells (ATCC) were cultured in Dulbecco's modified Eagle medium (DMEM, phenol red-free; Life Technologies) supplemented with 10% (v/v) fetal bovine serum (Life Technologies), 1 mM GlutaMAX (Life Technologies) and maintained at 37 °C in a humidified 5% (v/v) CO2 environment. The cells have integrated a histone H2B–HaloTag expressing plasmid via the piggyback transposase, under the selection of 500 μg/ml Geneticin (Life Technologies). This ‘H2B–Halo’ cell line undergoes regular mycoplasma testing by the Janelia Cell Culture Facility. Cells were imaged on a confocal microscope in the Janelia Imaging Facility (Zeiss 710, W Plan APO 20×/1.8 D) using the indicated filter sets.

### Comparison of JF_549_ and JF_525_

For the dye loading comparison (**Fig 1h**), H2B-Halo cells were stained for varying amounts of time with 100 nM of either JF_525_–HaloTag ligand 14 or JF_549_–HaloTag ligand **15**. The dye was washed from the cells and subsequently labeled with JF_646_–HaloTag ligand **4** at 1 μM for 30 min. Fluorescence of JF_646_–HaloTag ligand was quantified from the nuclear signals in summed confocal image stacks collected with 633 nm Ex/638-759 nm Em and analysed using Fiji.^45^ The integrated density of the nuclear signal was corrected by subtracting the integrated density of adjacent background regions. Labeling is expressed as the percent of the JF_646_–HaloTag fluorescence displaced by the JF_525_– and JF_549_–HaloTag ligands. The nuclear staining of these cells by the JF_525_–HaloTag ligand (**Fig.1g**) is displayed as a maximum intensity projection of confocal image stacks, 514 nm Ex/530-657 Em.

### Comparison of JF_503_ and other 488 nm-excited dyes

H2B-Halo COS7 cells were labeled with 200 nM of HaloTag^®^ R110Direct^™^ ligand (**19**, Promega), HaloTag ligand **20**^26^, or JF_503_–HaloTag ligand **18** over a time course of 0–;2 h. Cells were washed 2× with PBS and fixed with 4% w/v paraformaldehyde in 0.1 M phosphate for 30 min, followed by two more washes with PBS. Cells were imaged using confocal microscopy with 488 nm Ex/515-565 nm Em. The nuclear staining of these cells by the JF_525_–HaloTag ligand (**Fig. 2c**) is displayed as a maximum intensity projection of confocal image stacks. Corrected nuclear fluorescence was calculated as above to determine the cell loading profile (**Supplementary Fig. 1c**). To test the relative bleaching rates of these three dyes under imaging conditions (**Fig. 2d**), cells were stained and fixed as previously described for 2 h and then bleached with 488 nm at twice the typical power and imaged after each of 70 cycles. Bleached fluorescence data are normalized to the initial fluorescence levels.

### Comparison of HaloTag ligands 4, 23, 24, 27, and 28 in cells

H2B–Halo COS7 cells were labeled with 250 nM of JF_608_–HaloTag ligand (**23**), JF_585_–HaloTag ligand (**24**), JF_646_-HaloTag ligand (**4**), JF_635_-HaloTag ligand (**27**), or SiTMR–HaloTag ligand (**28**; **Supplementary Fig.1d**) and imaged by confocal microscopy using 594 nm Ex/599-734 nm Em (JF_608_ and JF_585_) or 633 nm Ex/638-759 nm Em (JF_646_ JF_635_, or SiTMR). All five samples were imaged via confocal microscopy without washing out the dyes. Signal to noise ratios were determined using the mean fluorescence of the nuclei relative to a region adjacent to each nuclei using Fiji^45^ (n = **152**–**275** areas as noted; **Fig.3g–j**, **Supplementary Fig.1f**).

### Staining with SNAP-tag ligands

COS7 cells were transfected with pSNAP-H2B (New England Biolabs) and stable integration of this plasmid was selected for using 600 μg/ml Geneticin^®^ (Life Technologies). This cell line expresses the histone H2B protein fused to the 26m version of the SNAP protein. Cells were stained with four different dyes as follows; JF_503_–cpSNAP-tag ligand (**29**, 2 μM for 90 min), JF525–SNAP-tag ligand (**30**, 3 μM for 30 min), JF_585_–SNAP-tag ligand (**31**, 2 μM for 3 hr with 0.2% w/v Pluronic F-127), JF_635_–SNAP-tag ligand (32, 2 μM for 2 h with 0.2% w/v Pluronic F-127). After staining, cells were washed three times with complete media, followed by a 20-min incubation in a 37 °C, 5% CO_2_, humidified incubator. The media was replaced again immediately prior to imaging.

### Staining of *Drosophila* larvae

The central nervous system of a third instar *Drosophila melanogaster* was dissected in physiological saline. This larva expressed myristoylated HaloTag in the ‘Basin’ neurons under control of the enhancer fragment R72F11.^46^ The isolated nervous system was incubated in physiological saline containing 1 μM JF_635_–HaloTag ligand (**27**) for 10 min at room temperature. The specimen was then embedded in agarose and imaged with the SiMView light-sheet microscope.^31^

### General information for mouse *in vivo* experiments

Male mice, 3–8 months old, were used for viral infection, dye injection, and *in vivo* imaging of neurons in the visual cortex (V1): The Scnn1a-Tg3-Cre (Jax no. 009613) line was used for imaging in L4; and Rbp4-Cre mice (MMRRC no. 031125-UCD) were used for imaging in L5 neurons. All experimental protocols were conducted according to the National Institutes of Health guidelines for animal research and were approved by the Institutional Animal Care and Use Committee at the Janelia Research Campus, HHMI.

### Cranial window implant and virus injection

A craniotomy was carried out at the same time as the virus injection to provide optical access for *in vivo* imaging experiments. Mice were anesthetized with isoflurane (1–2% v/v in O_2_) and given the analgesic buprenorphine (SC, 0.3 mg/kg). Using aseptic technique, a 3.5 mm-diameter craniotomy was made over the left V1 region of the brain of anaesthetized mouse (center: 3.4 mm posterior to Bregma; 2.7 mm lateral from midline). The dura was left intact. HaloTag and GCaMP6s was cotranduced using the viral vector: AAV2/1.synapsin.FLEX.GCaMP6s.P2A.HaloTag.WPRE (∼5 × 10^12^ infectious units per ml, 30 nl per site). The virus was injected using a glass pipette beveled at 45° with a 15–20-μm opening and back-filled with mineral oil. A fitted plunger controlled by a hydraulic manipulator (Narashige, MO10) was inserted into the pipette and used to load and inject the solution into 6 sites of left V1 (3.4–4.4 mm posterior to Bregma; 2.2–2.8 mm lateral from midline; ∼0.5 mm distance between each injection site, 0.5 mm below pia). A cranial window made of a single glass coverslip (Fisher Scientific no. 1.5) was embedded in the craniotomy and sealed in place with dental acrylic. A titanium head-post was attached to the skull with cyanoacrylate glue and dental acrylic.

### Dye Administration

JF_585_–HaloTag ligand (**24**) was administered to mice 3-4 weeks after the cranial window installation and viral injection. Dye solution was prepared by first dissolving 100 nmol (76 μg) of JF_585_–HaloTag ligand (**24**) in 20 μL DMSO. After vortexing, 20 μL of a Pluronic F-127 solution (20% w/w in DMSO) was added and this stock solution was diluted into 100 μL or 200 μL sterile saline for IV (tail vein) or IP injection, respectively.

### *In vivo* wide-field imaging and analysis

Mice were head-fixed and awake during the imaging period and were therefor habituated to experimental handling and head fixation starting 1-week post-surgery. During each habituation session, mice were head-fixed onto the sample stage with body restrained under a half-cylindrical cover. The habituation procedure was repeated 3–4 times for each animal for a duration of 15–60 minutes. For *in vivo* wide-field imaging, an external fluorescence light source (Leica EL6000, Leica) was used for excitation of GCaMP6s (green channel) and JF_585_–HaloTag ligand (red channel). Images were acquired via Leica Application Suite 4.5 (Leica). Wide-field images in green (1 second exposure) and red (4 second exposure) channels were acquired at multiple time intervals over two weeks under the same imaging conditions and the images were aligned with the Stackreg plugin in ImageJ. The mean values in the same area of red and green channels were plotted to track the labeling kinetics and turnover of JF_585_–HaloTag *in vivo*.

### *In vivo* two-photon imaging and analysis

For *in vivo* two-photon imaging, GCaMP6s and JF_585_–HaloTag were excited at 940 nm and 1100 nm, respectively, using a femtosecond laser source (InSight DeepSee, Spectra-Physics), and imaged using an Olympus 25 × 1.05 NA objective and a homebuilt two-photon microscope^47^. Images were acquired from 200 to 550 μm below the pia with post-objective power ranging between 20 and 60 mW. No photobleaching or photodamage of tissue was observed. Typical imaging settings were composed of 256 × 256 pixels, with 1.2 μm per pixel, and a ∼3 Hz frame rate. The time-lapse calcium images of spontaneous neuronal activity in awake, head fixed mice were recorded and analyzed with custom programs written in MATLAB (Mathworks). Lateral motion present in head-fixed awake mice was corrected using a cross-correlation-based registration algorithm^48^, where cross-correlation was calculated to determine frame shift in x and y directions. Cortical neurons were outlined by hand as regions of interest (ROIs). The fluorescence time course of each ROI was used to calculate its calcium transient as ΔF/F (%) = (F−F_0_)/F_0_ × 100, with the baseline fluorescence F_0_ being the mode of the fluorescence intensity histogram of this ROI.

**Supplementary Figure 1.**
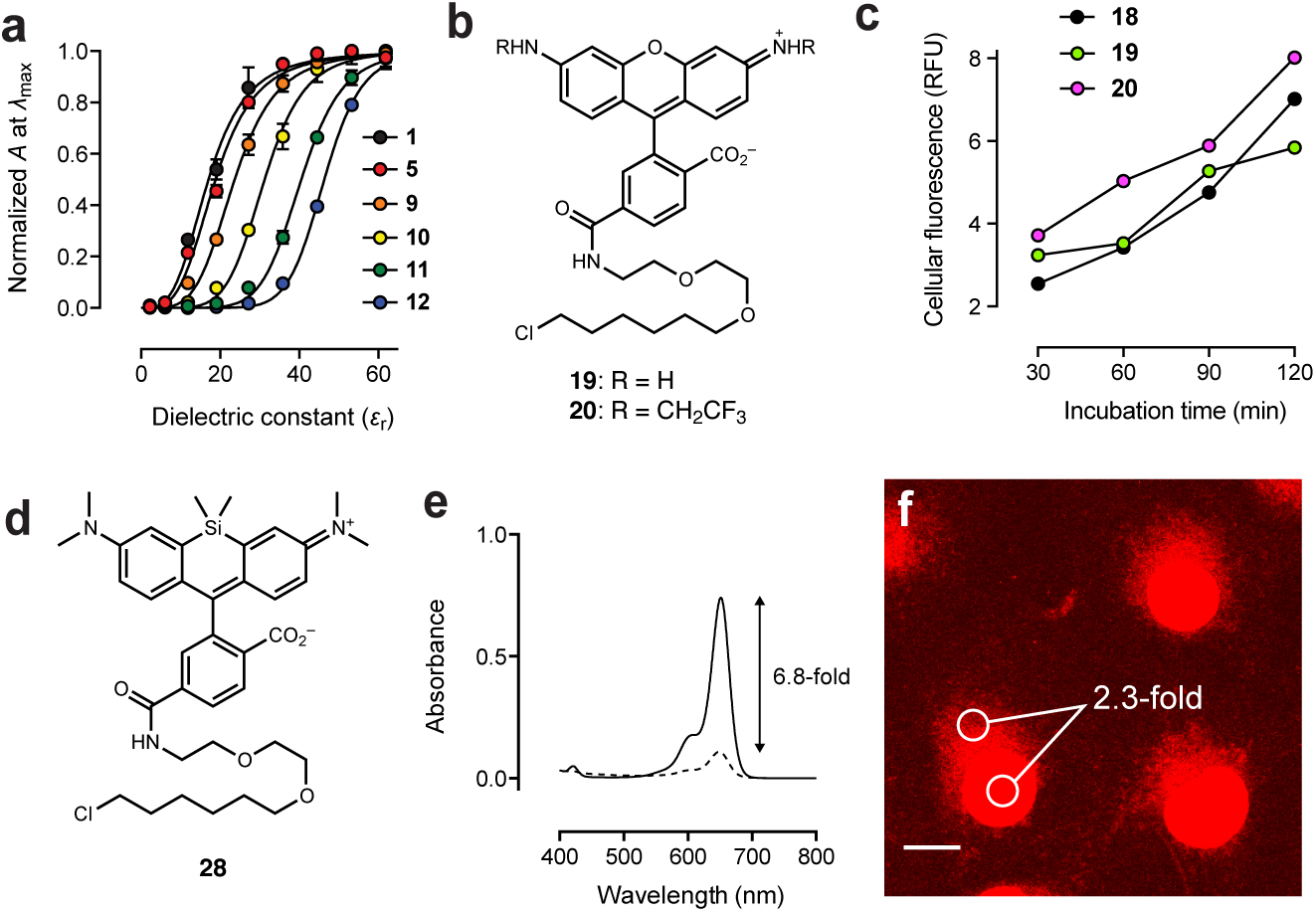
(**a**) Normalized absorption vs. dielectric constant (*ε_r_*) for dyes **1**, **5**, and **9**–**12**. (**b**) Chemical structures of known HaloTag ligands **19** and **20**. (**c**) Plot of average cellular fluorescence *vs*. incubation time for live cells loaded with ligands **18**–**20**. (**d**) Chemical structure of SiTMR–HaloTag ligand **28**. (**e**) Absorbance of SiTMR–HaloTag ligand **28** in the absence (−HT) or presence (+HT) of excess HaloTag protein. (**f**) Images of COS7 cells expressing HaloTag–histone H2B fusion and labeled with 250 nM of HaloTag ligand 28 for 1 h and imaged directly without washing. The number indicates mean signal (**nuclear**) to background (cytosol; SNB) in three fields of view (n = 152 areas). This image was taken with identical microscope settings to those used with ligands **4** and **27** (**Fig. 3i–j**). Scale bar: 15 μm.

**Supplementary Figure 2.**
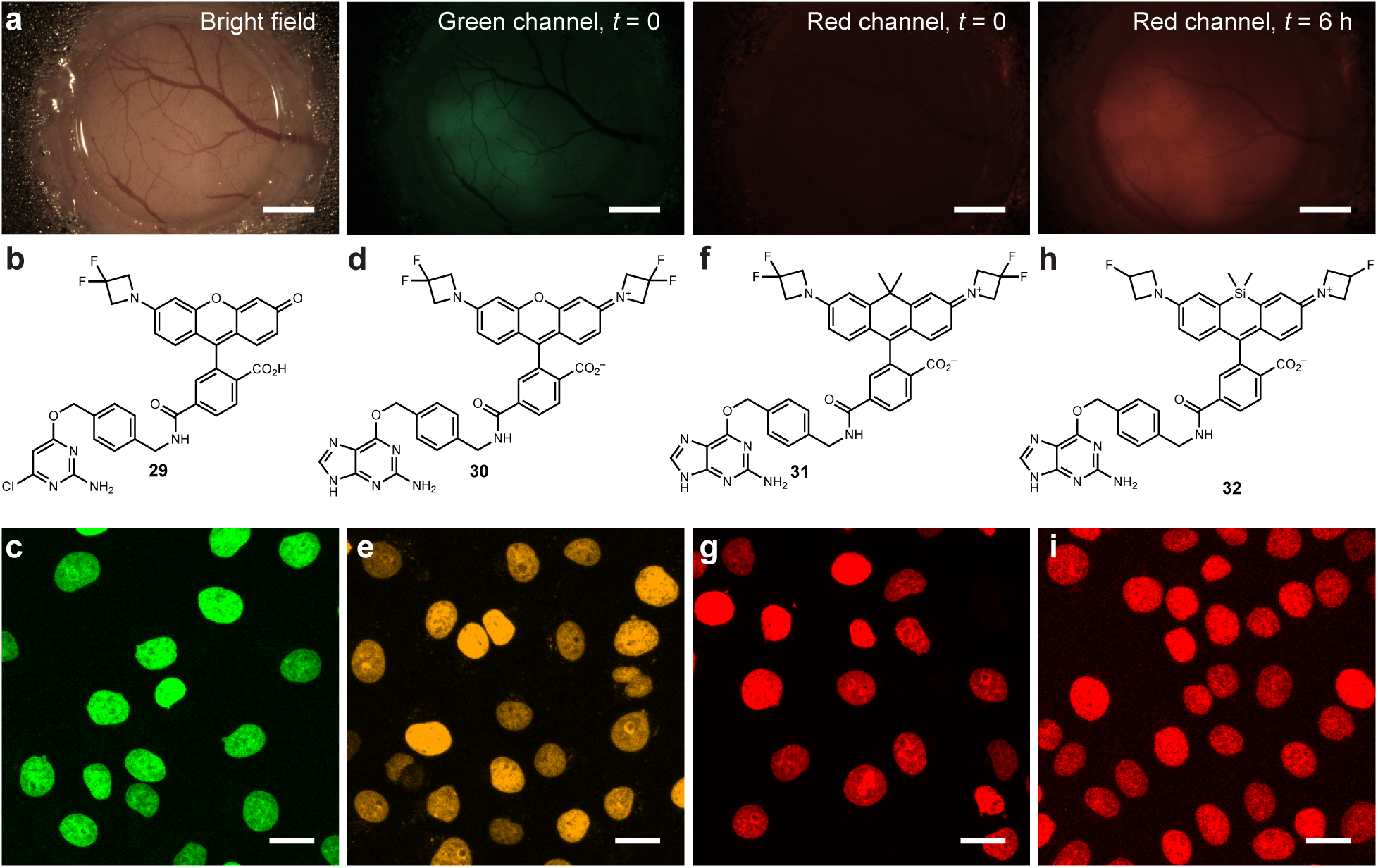
(**a**) Representative images from the labeling time course for JF_585_–HaloTag ligand (**24**) *in vivo*. Bright field image showing cranial window and epi-fluorescence images of green (GCaMP6; *t* = 0) and red (JF_585_, *t* = 0 and 6 h). (**b,c**) Structure of JF_503_–cpSNAP–tag ligand (**29**) and image of live COS7 cells expressing SNAP-tag–histone H2B and stained with ligand **29**. (**d,e**) Structure of JF_525_–SNAP-tag ligand (**30**) and image of live COS7 cells expressing SNAP-tag–histone H2B and stained with ligand **30**. (**f,g**) Structure of JF_585_–SNAP-tag ligand (**31**) and image of live COS7 cells expressing SNAP-tag-histone H2B and stained with ligand **31**. (**h,i**) Structure of JF_635_–SNAP-tag ligand (**32**) and image of live COS7 cells expressing SNAP-tag–histone H2B and stained with ligand **32**. Scale bars for images in **c**, **e**, **g**, and **i**: 15 μm.

#### Supplementary Video 1

Layer 5 neurons expressing GCaMP6s and HaloTag were labeled with JF_585_–HaloTag ligand (**24**) through intraperitoneal (IP) injection and imaged with two-photon fluorescence microscopy. JF_585_ was excited at 1100 nm and the stack (307 μm × 307 μm × 530 μm) was acquired from 50 to 580 μm below dura mater at 2 μm step in Z. 3D movie was made by the ImageJ 3D view plugin (unit is in μm).

